# Comparison of chronic aerosol inhalation with combustible cigarette and e-cigarette on the psychiatric behaviors and neuroimmune profile in mice

**DOI:** 10.1101/2023.12.14.571615

**Authors:** Zhibin Xu, Jiayan Ren, Xiaoyuan Jing, Zhi-zhun Mo, Zixuan Li, Yiqing Zhao, Ruoxi Wang, Zehong Wu, Xin-tao Jiang, Ye Tian, Liping Wang, Zuxin Chen, Xin-an Liu

## Abstract

With the worldwide use of electronic cigarettes (e-cigarettes) as a substitution for tobacco, the effects of e-cigarette vapor exposure on human health have been investigated. However, the comparison of long-term effects of aerosol inhalation with combustible cigarette and e-cigarette on the psychiatric behaviors has not been fully revealed. The present study examines the distinct effects of combustible cigarette and e-cigarette on anxiety, depression, cognition, and social behaviors. Here we found that the combustible cigarette induced the higher level of anxiety after long-term inhalation compared to the e-cigarettes with or without the tobacco flavor. Since the mechanism of action on the psychiatric behaviors entails the alterations on the neuroimmune-sensors and principal regulators such as glial cells, we further profiled the alterations of microglia and astrocytes by chronic inhalation of combustible tobacco cigarette, specifically the negative correlations between the IBA-1 level in the locus coeruleus (LC) and the latency to nest in VLT; as well as the GFAP level in LC and the open arm time in EPM were observed. Our current data provided an insight into the less impact of e-cigarettes on the anxiety-like behaviors and neuroimmune activation compared to combustible tobacco cigarettes which is not related to the flavor in e-cigarette, and the modulation on the neuroimmune signals in LC could be a therapeutic target for smoking-related anxiety.

## 1. Introduction

Each year, tobacco use directly causes the deaths of more than 8 million individuals globally [1]. Strategies to cut down on tobacco usage are therefore urgently needed. Electronic cigarettes (or "e-cigarettes"), commonly referred to as "electronic nicotine delivery systems" (ENDS), have gained popularity as a nicotine delivery product in place of conventional combustible cigarettes in recent years. E-cigarette use has grown significantly in recent years, a tendency that grew even faster during the COVID-19 pandemic [2]. The retail e-cigarette sector is growing on a global scale. However, perspectives on the health benefits of e-cigarettes vary, especially given that these devices have become more popular among teenagers and young adults even while consumption of traditional tobacco products has consistently decreased in this age group over time [3].

Many evidence has been obtained from cell cultures [4–6], animal models [7–9] and human subjects [10–13] to compare the vaping with smoking and assesses the impact of e-cigarettes on human health, especially the lung functions as well as cancer, oxidative stress, and inflammation related molecular signals [14–19]. A few studies on the cognitive and affective alterations after long-term exposure to tobacco smoke or e-cigarette vapour have demonstrated that, e-cigarette nicotine vapor exposure increased levels of locomotor activity during late prenatal and early postnatal life [20]; It has been recently reported that nicotine vapor exposure impairs the risk-based decision making in adult rats [21]; our previous work indicated that menthol flavor in e-cigarette vapor increases social behaviors in mice [22]. The potential mechanisms may include nicotine-related alterations of neurotransmitters and chemicals that signal inflammation in certain brain regions, such as the decreased dopamine concentration in the striatum and GABA in the frontal cortex, while increased levels of glutamate in the striatum and glutamine in the frontal cortex and striatum by chronic inhalation of e-cigarette vapor containing nicotine [23]; the reduced gene expression of *Ngfr* and *Bdnf*, and increased expression of IBA-1, a specific marker of microglia in the hippocampus [24]; changes on the CNS development-related gene profile in the frontal cortex [25]; and cytokines such as interleukin IL-4 and interferon-gamma (IFN-γ) in the diencephalon, as well as levels of hippocampal IFN-γ, and IL-6 in the cerebellum in the offspring of mice exposed prenatally to e-cigarette aerosols [26]. The reductions on the *CHRNB2* in the nucleus accumbens and the medial prefrontal cortex (mPFC) [21], and increased α-7 nicotinic acetylcholine receptor in the frontal cortex (FC) and striatum (STR) [27], were observed in animal models administrated with chronic nicotine vapor.

However, it is unclear how the immune system in the brain affects the expression of nicotinic receptors as well as the activities of the related neural circuits to mediate alterations in cognition and psychiatric behaviors. Further, understanding the physiological differences between e-cigarettes and tobacco, particularly in terms of neuropsychiatric effects, is important since e-cigarettes are by far the most popular tobacco substitute [28–30], the amount of study in this field is, however, insufficient. It has been reported that, e-cigarette exposure exhibits less toxic effects in lungs compared to cigarette smoke [31–33]; while exposure to e-cigarette and combustible tobacco cigarette may induce similar cardiac alterations in rats regarding the myocardial oxidative stress and inflammation [34]. Additionally, flavors in the e-cigarettes have also been shown to cause neuroinflammation which may contributes to behavioral changes at varying levels [35]. Thus we knew very little on the competitive effects of exposure to electronic and combustible cigarette aerosols regarding the investigations on the cognition and psychiatric effects, as well as the correlated alterations in the central nervous system. Here we administrated the mice with long-term exposure to tobacco flavored e-cigarette and combustible tobacco, and broadly assessed their roles on psychiatric behaviors. The activation of brain immune system as well as neuroinflammation were also evaluated. Our current work depicted the Comparative atlas on the profile of psychiatric behaviors and neuroimmune in mice with chronic exposure of combustible cigarette and tobacco-flavored e-cigarette for the first time, which will further allow us to understand the distinct neurophysiological effects between e-cigarettes and tobacco.

## 2. Experimental procedures

### 2.1. Animals

Two-month-old male C57BL/6J mice (Guangdong Gempharmatech Co., Ltd) were housed in cages in groups of four mice, lived in a 20 °C∼26 °C temperature and 40 %∼ 70 % humidity animal facility on a 12 h/12 h light/dark cycle (lights on at 8 a.m. and off at 8 p.m.), with ad libitum access to food and water. The bedding consisted of wood shavings was updated weekly to keep a comfortable environment. A total number of forty mice were assigned averagely and randomly to five groups. All animal experiments and procedures were respected the animal welfare guideline to minimize the number of mice used and their suffering, and approved by the Animal Care and Use Ethics Committee of the Shenzhen Institutes of Advanced Technology, Chinese Academy of Sciences.

### 2.2. E-Cigarette exposure procedures

After quarantine and acclimatization to environment for one week, every eight mice were sent to expose in a single part of 5 L volume acrylic whole-body chamber which allowed them to explore freely in one single session. The in-Expose e-cigarette device (SCIREQ Scientific Respiratory Equipment Inc.) was used to connect the chamber to accurately provide the vapor and fresh air for mice inhalation, and also expelled exhaust gas. The exposure duration was set up as 30 minutes per session per day for twenty weeks. The bias flow rate was 2 L/min, and the device generated 2 puffs/min and one puff delivered 55 ml volume vapor. Five groups were respectively referred to expose as follows: (1) Veh cont: Vehicle control of 50 % propylene glycol (PG) + 50 % vegetable glycerin (VG); (2) Nico: 4 % Nicotine + 46 % PG + 50 % VG; (3) Nico + TF: nicotine with tobacco flavoring: 10 % of tobacco flavoring agent + 4 % of Nicotine + 35 % of PG + 51 % of VG; (4) Tobacco: Hongtashan Classic 1956 Hard Cigarette contained 1.1 mg nicotine. The concentration of nicotine in the aerosol of each group was adjusted as shown as Table S1.

### 2.3. Behavioral Assessment

#### 2.3.1 Conditional place preference test

An unbiased conditioned place preference test (CPP) paradigm was used in this study. In brief, place conditioning chambers consisted of two distinct compartments (40 cm length × 60 cm width × 40 cm height) with different geometric patterns, separated by a cross baffle that allowed access to either side of the chamber. In the preconditioning phase (before e-cigarette aerosols exposure), animals were allowed to move freely between compartments for 10 min once daily for 3 days. Moving trajectory and time spent in each compartment was recorded as baseline measurements. These data were used to separate the animals into groups of approximately equal bias. The last 7 days in model establish phase were the conditioning phase, during which all groups received e-cigarette aerosols exposure respectively and then were immediately confined in the same side of compartment for 10 min. 4 hours later animals were confined in the opposite compartment for 10 min. After the last exposure, animals were immediately placed in a CPP compartments without cross baffle for 10 min. Moving trajectory and time spent in each compartment was recorded for subsequent analysis.

#### 2.3.2 Elevated plus maze

The elevated plus maze (EPM) test was widely designed to characterize anxiety levels of rodent which applied drugs and new chemical, since it has the advantage of unpainful and non-invasive operation. The EPM apparatus has two pair elevated arms, one pair open (25 cm length × 5 cm width × 15 cm height) and one pair closed (25 cm length × 5 cm width × 15 cm height) cross radiated from a central open zone. As soon as the experiment started, the animal was placed on the center zone facing an open arm, and the overhead camera was used to record 5-min activities of the animal. The time the animal spent in either the open or closed arms and their immobility time was recorded, which reflected the anxiety level of the animal.

#### 2.3.3 Hot plate test

The hot plate test is a simple way to measure the basal responsiveness under nociceptive stimulation. A heat source on a 55 ± 1 °C hot surface plays a role of nociceptive stimulation in the hot plate test. Basically, when the mice touch the hot surface, they will show an aversive response by licking a hind paw, vocalization, or jumping off the plate. The time between the touch to the thermal stimulation and the first overt aversive response was recorded as a nociceptive threshold in this test and the mouse was quickly taken away from the hot plate immediately after responding or after a maximum of 30 s (cut-off), to prevent tissue damage.

#### 2.3.4 Tail suspension test

The tail suspension test (TST) is mainly used to induce mice behavioral despair states which similar to depression. The mice would escape the stressful situation by instinct when suspended in the air. After a period of time, the mice ceased to struggle and gradually became subtle as immobility. The mice hanging without any limb or body movement except passive swinging was regarded as immobility. During the 6-min test, the activities of the mice which were separately and without interaction with each other, recorded by an automatic measurement device.

#### 2.3.5 Visual looming test

The visual looming test was carried out in a closed acrylic box (40 cm length × 40 cm width × 30 cm height) which placed a sheltered nest in the corner. An LCD display was hanged on the ceiling to present multiple upper field looming stimulus, which was a black dot expanding from a visual angle of 2 to 20 degree in a timing of 0.3 s with the speed of 60 degree per second. In a total time of 4.5 s the expanding action was repeated 15 times. The interval of action was 0.066 s. The movement trajectory of mice was recorded by an overhead camera. The onset time and duration of the mice expressed escape behavior and the duration hiding in the nest were recorded. The mice were first allowed to freely explore in a 5-min session.

#### 2.3.6 Y maze spontaneous alternation test

The Y maze test was routinely conducted to evaluate spatial working memory using a symmetrical three-arm apparatus. The dimensions of each arm were (42 cm length × 4 cm width × 25 cm height). The mice were allowed to explore freely all three arms in a single trial of 5 min. Spontaneous alternation was defined as a triplet of sequential entries into three different arms. The percentage alternation was calculated by dividing spontaneous alternation by (the total number of alternations minus two).

#### 2.3.7 Three-chamber social test

The three-chamber social test is broadly used to assess cognition by observe general sociability and interest in social novelty in rodent. Testing occurs in a three-chamber acrylic box (42 cm length × 60 cm width × 25 cm height), in which each chamber (42 cm length × 20 cm width × 25 cm height) has a door leaving the mice can freely exploring. Before the test, the mice were placed in the corner of the center chamber to habituate the apparatus and two empty cups for 10 min. In session 1, stranger 1 was placed inside the cup located in top half center of left chamber while right chamber leaving an empty cup in bottom half center. Then the subject mouse was placed at the center of middle chamber. In session 2, stranger 2 was placed inside the empty cup mentioned before, while other operations remained unchanged. Each session was monitored for 10 min and using 70 % ethanol to eliminate olfactory cues. Time around and the number of interactions with each cup, and the time spent in each chamber was recorded. Calculations of preference of index and preference of chambers were described in previous studies[22].

### 2.4 Neurochemical studies

#### 2.4.1 Brain tissue harvesting

Twenty-four hours after last exposure, the mice were sacrificed. The brain was either rapidly perfused nor immediately dissected, inserted in 1.5 ml EP tubes, and quickly frozen on dry ice before being stored at stored to −80 □.

#### 2.4.2 Tissue processing for immuno-stain

The perfused brain was removed from the skull and postfixed in 4 % PFA at 4 □ for 24 h. Then put in 30 % sucrose solution for 48 h, absorb the surface water, and embed the brain with OCT at −20 □. Brains after postfixing were cryoprotected in 30 % sucrose in 1 × PBS for 48 h, absorb the surface water and thereafter embed with OCT at −20 □ Cryosectioning. The brains were sectioned (40 µm thickness) using a vibratome (Leica VT1000, Leica Microsystems). The areas of the brain that we focused on were the Locus coeruleus (LC), the ventral tegmental area (VTA), the hippocampus (HIP), the Substantia nigra (SNr), the prefrontal cortex (PFC), and the basolateral amygdala (BLA). Cryosectioned sections were washed three times in 1 × PBS to remove OCT. Blocked with 10 % normal goat serum in 0.3 % PBST for 1.5 h at room temperature, and thereafter incubated with primary antibodies in 0.1 % PBST+5 %NGS at 4 □ overnight. The sections were thereafter washed three times in 1 × PBS and incubated with a species-specific fluorophore conjugated secondary antibody in 1 × PBS for 2 h at room temperature. After secondary antibody incubation, dilute DAPI stock solution (2 mg/ml) with PBS to prepare 1:5000 DAPI dilution, incubate at room temperature for 5 min. The sections were thereafter consecutively washed with 1 × PBS three times (10 min each). Cut sections were mounted on glass slides. All sections were coverslipped (Thermo Scientific) using anti-quenching agent.

#### 2.4.3 Histological mapping and quantification of microglia and astrocytes

Tiled whole-brain images were acquired of every second section at magnification × 10 using an Olympus bx61vs fluorescent microscope. Select images by overlapping a coverage of a brain atlas picture on the matched image in PS. Then, manual quantification in PS by marking target neurons with counting function.

#### 2.4.4 Inflammatory Cytokine Assay

The brain levels of 6 common inflammatory cytokines in Veh cont group, Nico group, Nico+TF group and Tobacco group were detected using an ABplex Mouse Th17 Secreted Cytokine 6-Plex Assay Kit (RK04293, ABclonal Technology, Wuhan, China). In short, the antibody molecules of different substances are covalently cross-linked to specific coded microspheres, and each coded microsphere corresponds to a corresponding detection item. Firstly, the fluorescence coding microspheres of different substances were mixed by liquid phase reaction, and then the compounds were added to react with the labeled fluorescein. Then, driven by the flowing sheath fluid, the microspheres pass continuously through red and green lasers. Finally, the fluorescence code of the microsphere was determined by red light, and the fluorescence intensity of the reporter molecule in the microsphere was determined by green light. The specific operation is as follows: adding 50 μL gradient standard or test samples per well, adding 5 μL microsphere suspension to incubate at 37 ℃ for 1 h, then washing once, adding 50 μL detection antibody solution to incubate at 37 ℃ for 0.5 h, washing once, adding 50 μL fluorescein solution to avoid light at 37 ℃ for 15 min, then adding 70 μL washing buffer, using ABplex-100 for detection.

### 2.5 Statistical analyses

The statistical data are described as mean values ± SEM. Two-way ANOVA followed by Tukey’s HSD test was used to compare each group as depicted in the figure legends. All data were calculated and plotted using the Prism 9.0 software (GraphPad Software Inc., San Diego, CA, United States).

## 3. Results

### 3.1 Chronic aerosol inhalation with combustible cigarette but not tobacco-flavored e-cigarette induces conditioned place aversion

E-cigarette products in the market are usually offer a very wide variety of flavoring agents mixed with nicotine, which is one of the biggest health concern for e-cigarettes [36, 37]. We here aim to evaluate whether the rewarding effects of ENDS can be modified by flavoring compound in e-cigarette. Menthol is one of the most prevalent and common flavors used in e-cigarettes, so we compared the behavioral responses in vapor exposed mice between nicotine alone group and group of nicotine mixed with menthol flavoring. To do this, adult male C57BL/6J mice were randomly assigned into three treatment groups (n=8 per group) which exposed to 1) propylene glycol and vegetable glycerol as a vehicle control (50:50, PG/VG, Veh cont); 2) PG/VG with 4 % (vol/vol) nicotine (Nico); 3) Vapor of 4 % nicotine with 10 % tobacco flavorings (Nico + TF). After exposure 43 days, a series of neuropsychiatric behaviors were evaluated between the above groups (Fig. 1, Fig. 2 and Fig. 3). Since we focused to investigate the merged effects of tobacco flavor in nicotine vapor, we here only use PG/VG as vehicle control in the following sets of behavioral assessments.

**Figure 1.**
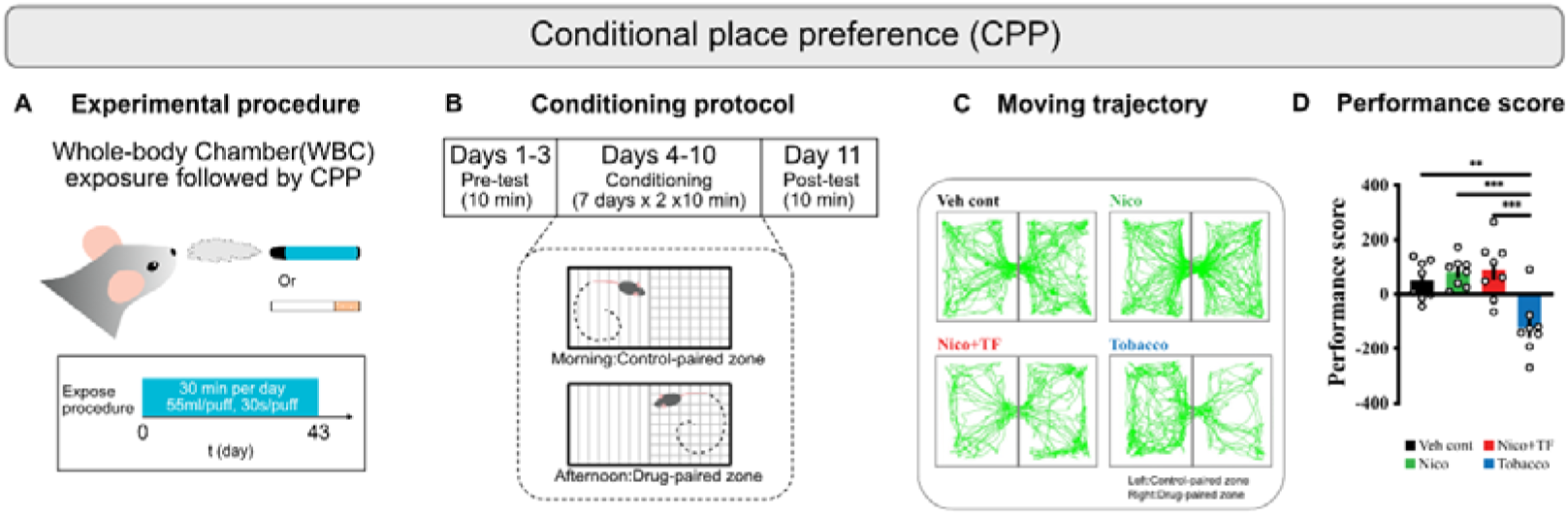
Overview of rewarding behavioral assessment in mice and the experimental workflow. (A) CPP was conducted after procedure of long-term whole-body chamber exposure. (B) schematic diagram of three stage in CPP test. (C) Moving trajectory by analyst from recording video. (D) A preference score in second (sec), which is compared by subtracting time spend in control-paired zone from time spend in drug-paired zone. \**p* < 0.05 as determined by Ordinary one-way ANOVA, multiple comparisons with every other group. Bars represent marginal means ± SEM. N=8 per group. Note: Veh cont, Vehicle control; Nico, Nicotine; Nico+TF, Nicotine with tobacco flavoring; Tobacco, Hongtashan Classic 1956 Hard Cigarette.

**Figure 2.**
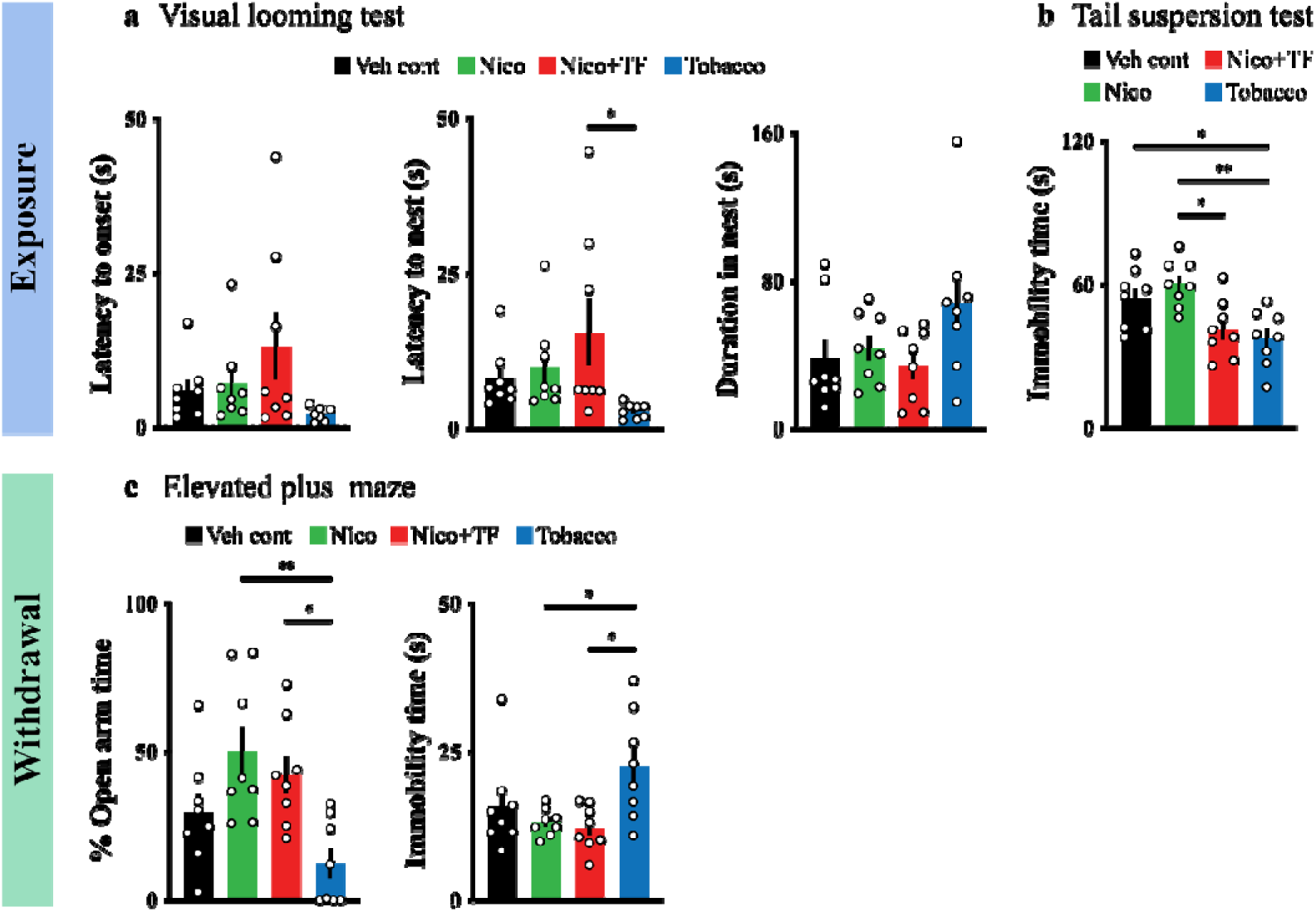
(a-b) Behavioral assessment without withdrawal. (a) The latency to onset behavior of mice during Visual looming test (sec). (b) Total immobility time of mice in 6-min suspension test. Withdrawal response assessments. (c) The immobility time of mice during the trial in EPM (sec). The percentage of time of staying in open arms to time of one trial ratio (%) and the immobility time of mice during the trial in EPM (s). \**p* < 0.05 and \*\**p* < 0.01 as determined by Ordinary one-way ANOVA, multiple comparisons with every other group. Bars represent marginal means ± SEM. N=8 per group. Note: Veh cont, Vehicle control; Nico, Nicotine; Nico+TF, Nicotine with tobacco flavoring; Tobacco, Hongtashan Classic 1956 Hard Cigarette. EPM, Elevated plus maze.

**Figure 3.**
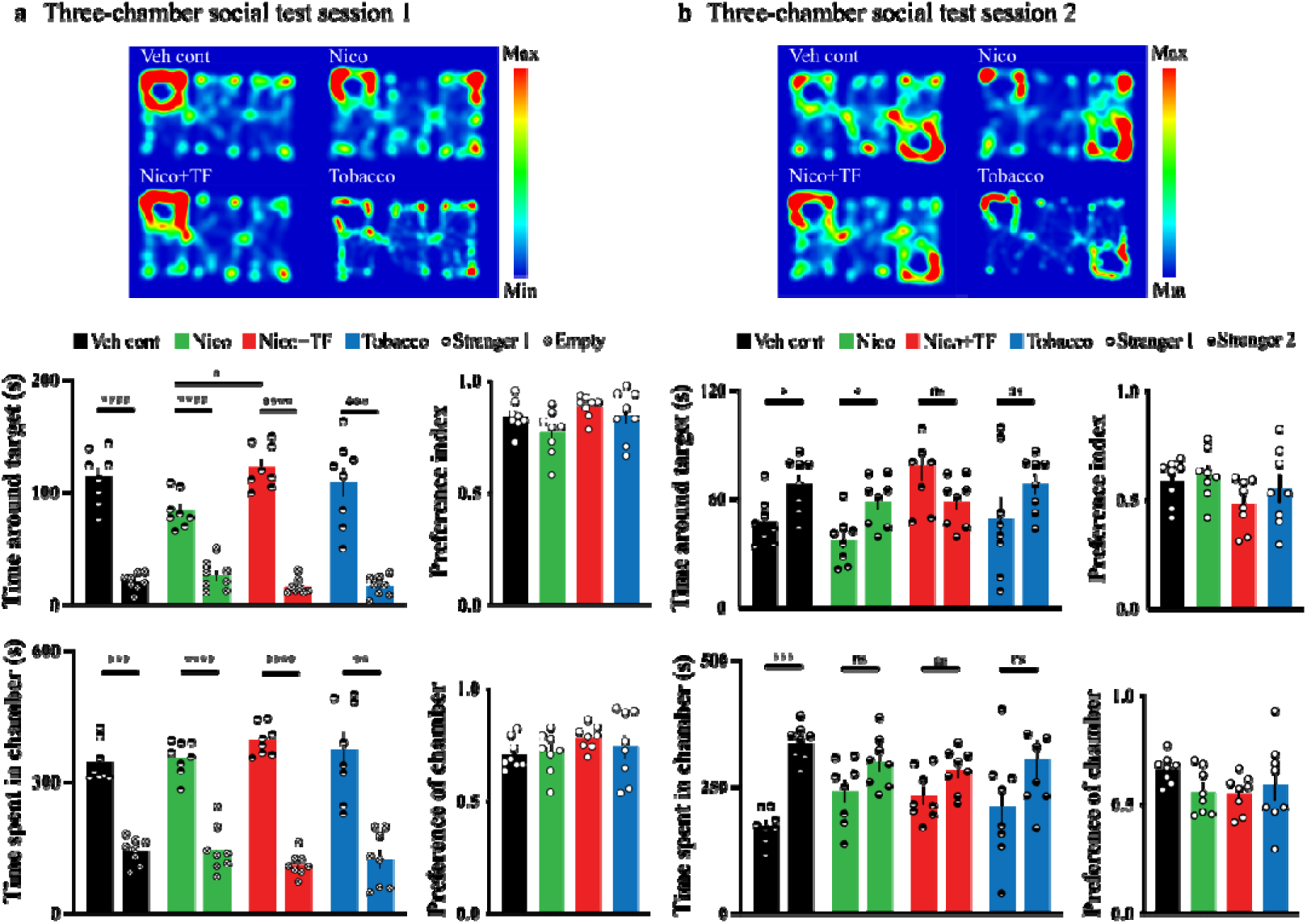
Social behavioral assessments in mice after long-term vapor exposure. Heatmaps: (a-b) represent the trajectory of free exposure in mice. Histogram: (a) Top left: time around stranger 1 mouse, top right: preference to contact with stranger 1 mouse; Bottom left: time spent in stranger 1 side chamber, bottom right: preference to stay in stranger 1 side chamber. (b) Top left: time around stranger 2 mouse, top right: preference to contact with stranger 2 mouse; Bottom left: time spent in stranger 2 side chamber, bottom right: preference to stay in stranger 2 side chamber. \**p* < 0.05, \*\**p* < 0.01, \*\*\**p* < 0.001, \*\*\*\**p* < 0.0001 as determined by Ordinary one-way ANOVA, multiple comparisons with every other group. Bars represent marginal means ± SEM. N=8 per group. Note: Veh cont, Vehicle control; Nico, Nicotine; Nico+TF, Nicotine with tobacco flavoring; Tobacco, Hongtashan Classic 1956 Hard Cigarette.

We compared the rewarding behavior in vapor exposed mice with nicotine, nicotine plus tobacco flavoring or tobacco. By performing the behavioral tests of CPP (with vapor exposure), we observed that long term vapor exposure of daily half-hour ENDS of nicotine with or without tobacco flavor did not cause “smoking addiction” in mice, but tobacco leading an aversive response (Fig. 1A). The value of preference score (Fig. 1D) revealed that there was not a significant difference on rewarding response between Veh control group and Nico group (*F*_(3, 28)_ = 10.55, *p* = 0.8922), as measured (50.41 ± 25.30 s) and (81.13 ± 19.15 s) respectively, moreover, with the same volumetric concentration (4 %) of nicotine, tobacco flavor-added group (Nico + TF) showed no change on the performance score compared with nicotine vapor itself (*F*_(3, 28)_ = 10.55, *p* = 0.9979), which indicated that tobacco flavor had no remarkable effect on addiction when mixed with nicotine in e-liquid. However, Tobacco group (−122.4 ± 36.54 s) showed a disgust response when compared with other groups, manifested a negative rewarding effect within tobacco product.

### 3.2 Chronic aerosol inhalation with combustible cigarette but not tobacco-flavored e-cigarette induces anxiety

We evaluated mood status in vapor exposed mice with nicotine, nicotine plus tobacco flavoring or tobacco. By performing the behavioral tests of EPM (after 48 hrs-withdrawal), the results of the ratio of staying in open arm time and immobility time showed no significant difference between Veh cont group and other groups, which suggested no significant withdrawal responses evaluated by anxiety-like behaviors after ENDS withdrawal (Fig. 2c). There was significant difference between Tobacco group and either Nico group (49.97 ± 8.500%, *F*_(3, 28)_ = 5.989, *p* = 0.0025) or Nico + TF group (42.30 ± 6.287%, *F*_(3, 28)_ = 5.989, *p* = 0.0190) in the ratio of staying in open arm time, and significant difference in immobility time between Tobacco group and either Nico group (13.14 ± 0.8091, *F*_(3, 28)_ = 4.188, *p* = 0.0333) and Nico + TF group (12.19 ± 1.347, *F*_(3, 28)_ = 4.188, *p* = 0.0166), suggested tobacco vapor making mice more stressful than nicotine or nicotine plus tobacco flavor vapor. Further, we investigated the depressive effects in TST (with vapor exposure) (Fig. 2b). We found significant difference in immobility time between Tobacco group and other groups, suggested tobacco has a depressive effect, while no significant difference between Veh cont group and either Nico group (*F*_(3, 28)_ = 6.600, *p* = 0.7138) or Nico + TF group (*F*_(3, 28)_ = 6.600, *p* = 0.1563).

### 3.3 No alterations on the social behaviors and spatial working memory after chronic aerosol inhalation with cigarette and e-cigarette

We observed no significant difference among all groups (*F* _(3,28)_ = 1.216, *p* = 0.3223, Supplemental Fig. S1-d) in Hot Plate Test (HPT). We also analyzed the innate response of them when faced a visual threat stimulation in Visual looming test (VLT) (Fig. 2a). We found no significant difference among all groups in latency to onset (*F* _(3,28)_ = 2.115, *p* = 0.1208), latency to nest (*F* _(3,28)_ = 2.829, *p* = 0.0565), duration in nest (*F* _(3,28)_ = 2.207, *p* = 0.1094), but there had a trend that tobacco group was lower than Nico group or Nico + TF group. Especially, tobacco group was significantly lower than Nico + TF group in indicator of latency to nest (*F* _(3,28)_ = 2.829, *p* = 0.0355), indicated tobacco has defect of innate response to a certain degree. We found significant difference among all groups in Y maze in two indicators (Supplemental Fig. S1-a, b, c). Firstly, there was no difference between Veh cont group and other groups in the percent of spontaneous alternation, but Nico + TF group was higher than other groups, specifically has significant difference with Nico group (*F* _(3,28)_ = 3.729, *p* = 0.0179), which indicated add tobacco flavor in nicotine could improve spatial learning performance to a certain degree (Supplemental Fig. S1-a). Secondly, compared with Veh cont group (*F* _(3,28)_ = 4.711, *p* = 0.0465) and Nico + TF group (*F* _(3,28)_ = 4.711, *p* = 0.0073), Tobacco group showed a significant decline in total entries (Supplemental Fig. S1-b), which illustrated tobacco could lower innate exploring driver. And total entries and total distance (Supplemental Fig. S1-b, c) to three arms of Nico + TF was the highest, which indicated tobacco flavor could drive mice more active to explore.

Social interaction is a highly conserved and complex neuropsychiatric behavior to safeguard survival [38]. Whether the additive of tobacco flavor into e-cigarette would change the social interaction was unknown. We evaluated the social behaviors of mice via three-chamber social tests in this study. Sociability was investigated in the sociability session of the test (Fig. 3a). Mice exposed to long-term ENDS with or without tobacco flavor showed normal sociability as assessed by interaction time and time spent in the target chamber. Mice from the Nico group appeared normal while having slightly less contact and socialization with the stranger mouse 1 than in the empty arena compared with that in the Veh cont group (No statistical significance, Fig. 3a histogram). Interestingly, the mice in the Nico + TF group spent more time (122.9 ± 6.933 s) to socialize with the stranger mouse 1 compared to the Nico group (83.67 ± 5.558 s, *F* _(3,28)_ = 3.422, *p* = 0.0240, Fig. 3a top left histogram) while the time spent in the chamber of stranger mouse 1 (*F* _(3,28)_ = 0.9030, *p* = 0.4521) and the preference of chamber was similar to the mice (*F* _(3,28)_ = 0.8932, *p* = 0.4569) of all the groups (Fig. 3a bottom histogram). In the test session of preference for social novelty (Fig. 3b), mice in Veh cont and Nico groups spent more time in contact and socializing with newly introduced strangers (stranger 2) than strangers 1, which is a normal manifestation of social memory and social novelty preference (*F* _(3,28)_ = 0.9597, *p* = 0.4254, Fig. 3b top left histogram), but there was no significant difference between Nico + TF and Tobacco groups. The Veh cont group spent more time in Stranger 2 room, while the other three groups did not differ significantly (Fig. 3b bottom left histogram), and there was no significant difference in chamber preference among the four groups (Fig. 3b bottom right histogram). Our data suggested that long exposure to tobacco-flavored ENDS may have compensatory enhancing effects on the sociability and preference for social novelty compared to the vapor exposure of nicotine alone.

### 3.4 Neuroimmune profile after chronic aerosol inhalation with cigarette and e-cigarette in mice

Neuroinflammation induces anxiety- and depressive-like behavior by modulating neuronal activity in the amygdala. Astrocytes are crucial regulators of innate and adaptive immune responses via affecting inflammatory process in the brain [39]. To explore the role of neuroimmune regulation in various neuropsychiatric behaviors. We tested cytokine levels in mice brains and immunofluorescence tests, and cytokine tests showed that the content of IL-17A in mouse brains in Nico + TF (*F*_(3,8)_ = 39.03, *p* = 0.0314, Fig. 4B) and Tobacco (*F*_(3,8)_ = 39.03, *p* = 0.0022, Fig. 4B) groups was significantly lower than that in Veh cont group. Compared with Nico group, IL-17A content in Nico + TF (*F*_(3,8)_ = 39.03, *p* < 0.0001, Fig. 4B) and Tobacco (*F*_(3,8)_ = 39.03, *p* < 0.0001, Fig. 4B) groups was also significantly decreased. Compared with Veh cont group, IL-17A content in Nico groups was also significantly increased (*F*_(3,8)_ = 39.03, *p* < 0.0001, Fig. 4B). The content of IL-1β in mice brain was significantly decreased in Nico + TF groups compared with Veh cont group (*F*_(3,8)_ = 17.13, *p* = 0.0047, Fig. 4C). Compared with Nico group, IL-1β content in Nico + TF (*F*_(3,8)_ = 17.13, *p* = 0.0007, Fig. 4C) and Tobacco (*F*_(3,8)_ = 17.13, *p* = 0.0153, Fig. 4C) group was significantly decreased. Compared with Nico group, TNF-α content in Tobacco group was significantly increased (*F*_(3,8)_ = 10.91, *p* = 0.0269, Fig. 4E), while IFN-γ content was significantly decreased (*F*_(3,8)_ = 5.281, *p* = 0.0494, Fig. 4F). Compared with Nico + TF group, TNF-α content in Veh cont (*F*_(3,8)_ = 10.91, *p* = 0.0141, Fig. 4E) and Tobacco (*F*_(3,8)_ = 10.91, *p* = 0.0047, Fig. 4E) group was significantly increased. Compared with Veh cont group, IFN-γ content in Tobacco group was significantly decreased (*F*_(3,8)_ = 5.281, *p* = 0.0300, Fig. 4F),There was no significant difference in IL-6 (*F*_(3,8)_ = 1.728, *p* = 0.2382, Fig. 4A) and IL-10 (*F*_(3,8)_ = 1.652, *p* = 0.2532, Fig. 4D) content among the four groups.

**Figure 4.**
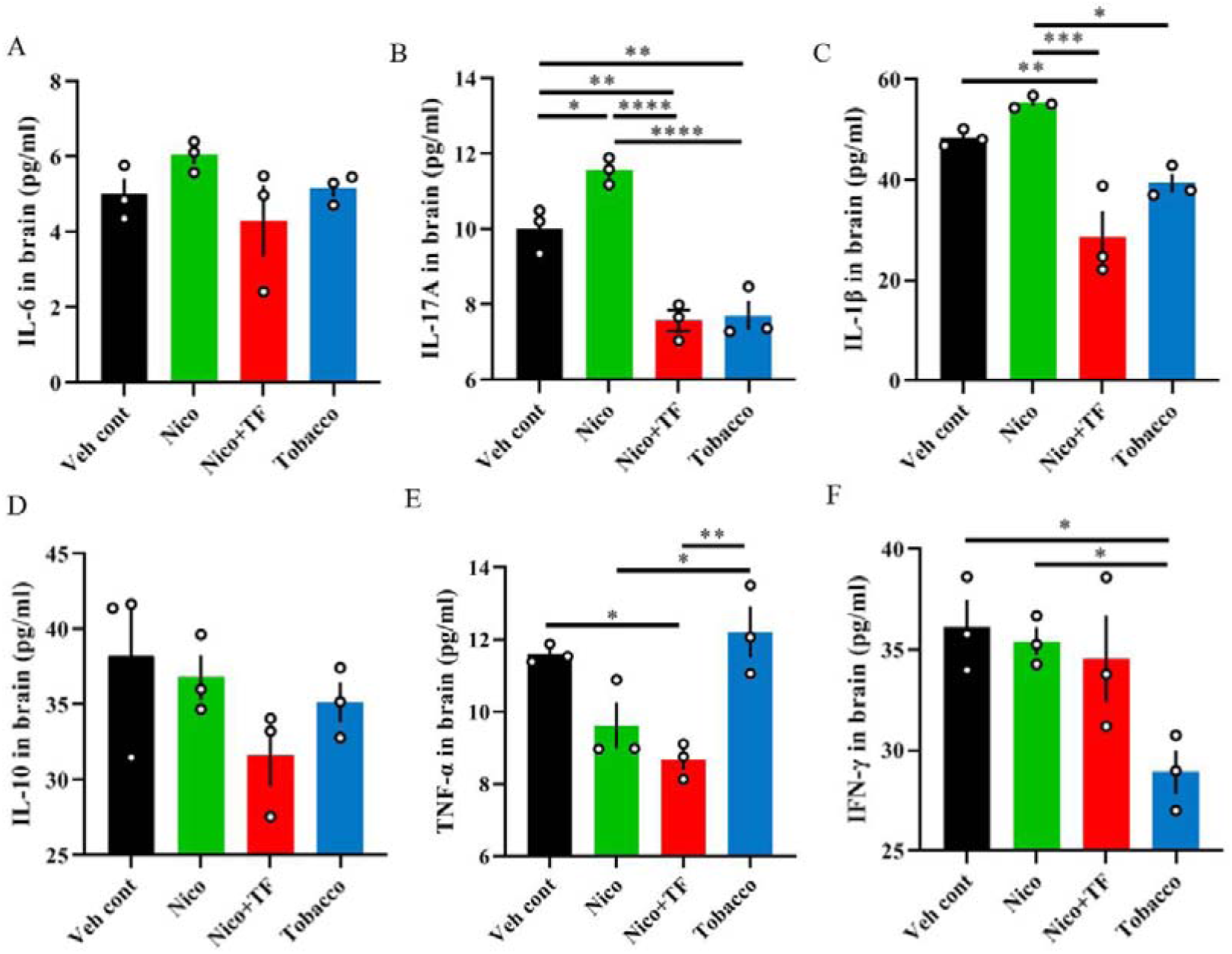
Results of brain cytokine profile by ELISA analyses. The levels of IL-6 (A), IL-17A (B), IL-1β (C), IL-10 (D), TNF-α (E), and IFN-γ (F) in the whole brain were detected using an ABplex Mouse Th17 Secreted Cytokine 6-Plex Assay Kit. \**p* < 0.05, \*\**p* < 0.01, \*\*\**p* < 0.001, \*\*\*\**p* < 0.0001 as determined by Ordinary one-way ANOVA, multiple comparisons with every other group. Bars represent marginal means ± SEM. N=3 per group. Note: Veh cont, Vehicle control; Nico, Nicotine; Nico+TF, Nicotine with tobacco flavoring; Tobacco, Hongtashan Classic 1956 Hard Cigarette.

Figure 5A shows the expression level of IBA-1 in brain regions of mice after different smoke exposure. Immunofluorescence results showed that the expression of IBA-1 in LC brain was significantly increased in Tobacco group compared with Nico group (*F*_(3,40)_ = 6.373, *p* = 0.0010, Fig. 5B) and Nico + TF group (*F*_(3,40)_ = 6.373, *p* = 0.0200, Fig. 5B). In VTA, Compared to the Tobacco group, the expression of IBA-1 was significantly lower than that in Veh cont group (*F*_(3,33)_ = 39.84, *p* < 0.0001, Fig. 5C), Nico group (*F*_(3,33)_ = 39.84, *p* < 0.0001, Fig. 5C) and Nico + TF (*F*_(3,33)_ = 39.84, *p* < 0.0001, Fig. 5C). In HIP, the expression of IBA-1 in Nico (*F*_(3,32)_ = 8.721, *p* = 0.0138, Fig. 5D), Nico + TF (*F*_(3,32)_ = 8.721, *p* = 0.0003, Fig. 5D) and Tobacco (*F*_(3,32)_ = 8.721, *p* = 0.0012, Fig. 5D) groups compared with Veh cont group was significantly increased. In SNr, compared with Veh cont group, the expression of IBA-1 was significantly increased in Nico + TF (*F*_(3,34)_ = 15.50, *p* = 0.0064, Fig. 5E) and Tobacco (*F*_(3,34)_ = 15.50, *p* < 0.0001, Fig. 5E). Compared with the Nico group, the expression of IBA-1 was significantly increased in Nico + TF (*F*_(3,34)_ = 15.50, *p* = 0.0085, Fig. 5E) and Tobacco (*F*_(3,34)_ = 15.50, *p* < 0.0001, Fig. 5E). However, there was no significant difference between the four groups in PFC (*F*_(3,32)_ = 2.784, *p* = 0.0567, Fig. 5F) and BLA (*F*_(3,32)_ = 1.553, *p* = 0.2198, Fig. 5G) brain regions.

**Figure 5.**
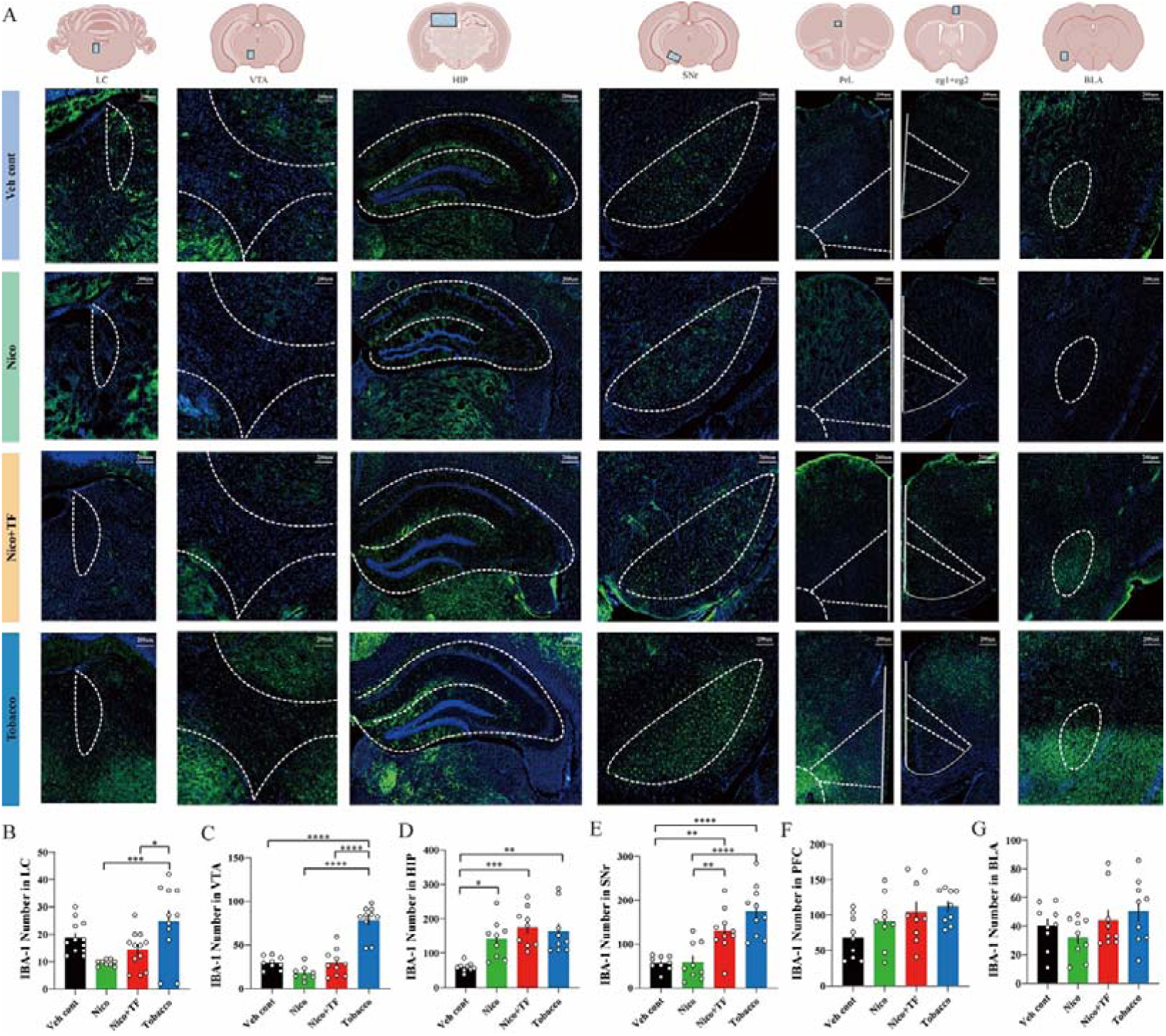
Effects of smoke exposure on IBA-1 expression in different brain regions. The schematic location and representative images of IBA-1 expression (A). Quantification of IBA-1 in LC (B), VTA (C), HIP (D), SNr (E), PFC (PrL+cg1+cg2) (F) and BLA (G). \**p* < 0.05, \*\**p* < 0.01, \*\*\**p* < 0.001, \*\*\*\**p* < 0.0001 as determined by Ordinary one-way ANOVA, multiple comparisons with every other group. Bars represent marginal means ± SEM. N=8-12 per group. Note: Veh cont, Vehicle control; Nico, Nicotine; Nico+TF, Nicotine with tobacco flavoring; Tobacco, Hongtashan Classic 1956 Hard Cigarette.

Figure 6A shows the expression level of GFAP in brain regions of mice after different smoke exposure. In LC brain area, the expression of GFAP in Tobacco group was significantly higher than that in Veh cont group (*F*_(3,40)_ = 7.984, *p* = 0.0149, Fig. 6B). Compared with Nico group, the expression of GFAP in Nico + TF (*F*_(3,40)_ = 7.984, *p* = 0.0292, Fig. 6B) and Tobacco group (*F*_(3,40)_ = 7.984, *p* = 0.0002, Fig. 6B) was significantly increased. In VTA, compared with Veh cont group, the expression of GFAP in Nico + TF group (*F*_(3,33)_ = 15.46, *p* < 0.0001, Fig. 6C) and Tobacco group (*F*_(3,33)_ = 15.46, *p* = 0.0192, Fig. 6C) was significantly increased. Compared with Nico group, the expression of GFAP in Nico + TF group (*F*_(3,33)_ = 15.46, *p* < 0.0001, Fig. 6C) and Tobacco group (*F*_(3,33)_ = 15.46, *p* = 0.0148, Fig. 6C) were significantly increased. In HIP, the expression of GFAP in Nico (*F*_(3,32)_ = 9.794, *p* = 0.0012, Fig. 6D) and Tobacco (*F*_(3,32)_ = 9.794, *p* = 0.0003, Fig. 6D) groups was significantly increased compared with Veh cont group. Compared with Nico + TF group, the expression of GFAP in Nico group (*F*_(3,32)_ = 9.794, *p* = 0.0420, Fig. 6D) and Tobacco group (*F*_(3,32)_ = 9.794, *p* = 0.0144, Fig. 6D) was also significantly increased. In SNr, compared with Veh cont group, the expression of GFAP in Nico + TF group (*F*_(3,34)_ = 8.078, *p* = 0.0079, Fig. 6E) and Tobacco group (*F*_(3,34)_ = 8.078, *p* = 0.0008, Fig. 6E) was significantly increased. Exposure to tobacco significantly increased the expression of GFAP compared with Nico (*F*_(3,34)_ = 8.078, *p* = 0.0132, Fig. 6E). In the brain region of PFC, the expression of GFAP in Veh cont group (*F*_(3,32)_ = 14.71, *p* < 0.0001, Fig. 6F), Nico + TF group (*F*_(3,32)_ = 14.71, *p* = 0.0003, Fig. 6F) and Tobacco group (*F*_(3,32)_ = 14.71, *p* = 0.0001, Fig. 6F) was significantly lower than that in Nico group. However, there was no significant difference between the four groups in BLA (*F*_(3,32)_ = 0.3770, *p* = 0.7702, Fig. 6G).

**Figure 6.**
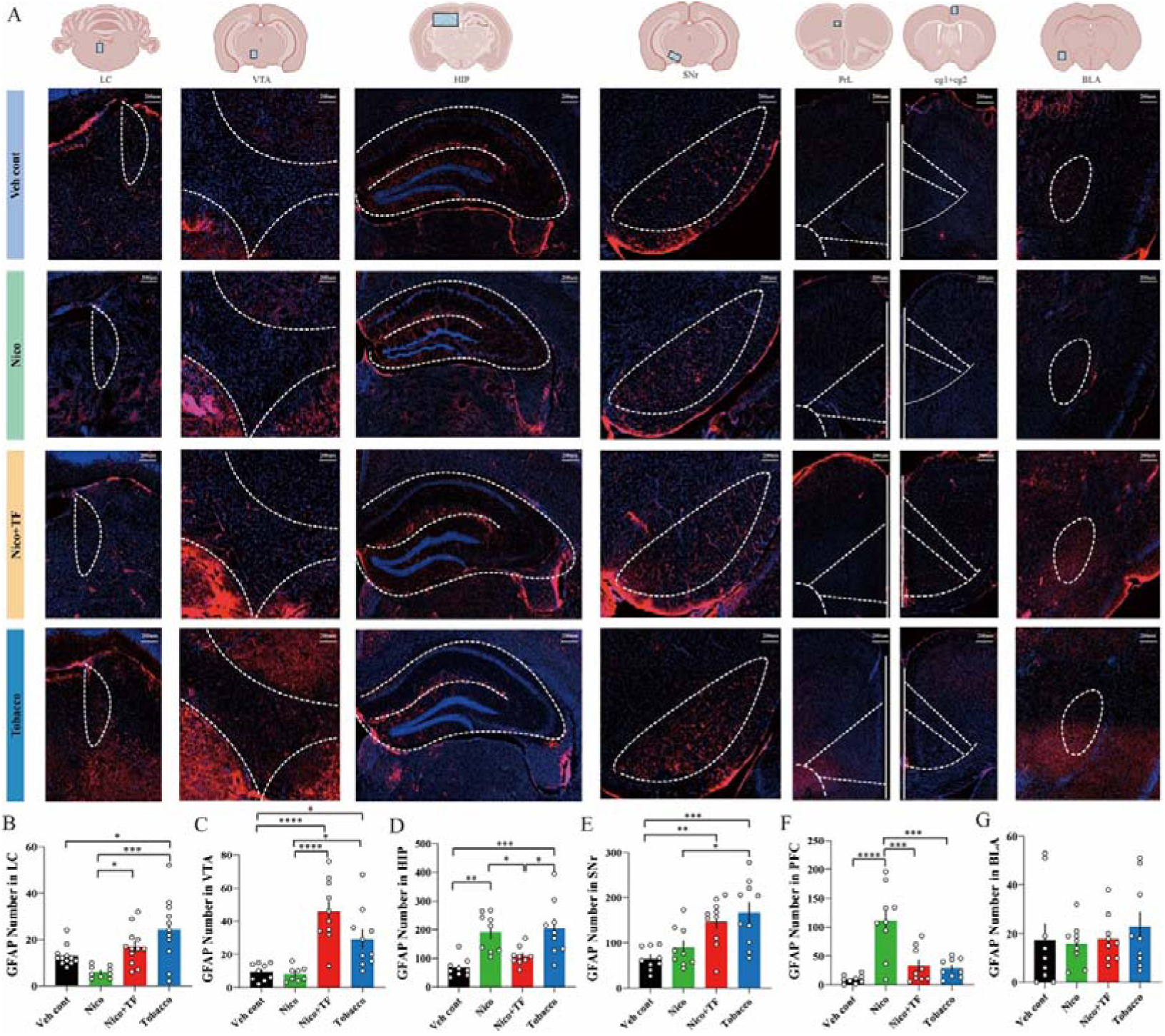
Effects of smoke exposure on GFPA expression in different brain regions. The schematic location and representative images of GFAP expression (A). Quantification of GFAP in LC (B), VTA (C), HIP (D), SNR (E), PFC (PrL+cg1+cg2) (F) and BLA (G). \**p* < 0.05, \*\**p* < 0.01, \*\*\**p* < 0.001, \*\*\*\**p* < 0.0001 as determined by Ordinary one-way ANOVA, multiple comparisons with every other group. Bars represent marginal means ± SEM. N=8-12 per group. Note: Veh cont, Vehicle control; Nico, Nicotine; Nico+TF, Nicotine with tobacco flavoring; Tobacco, Hongtashan Classic 1956 Hard Cigarette.

To explore the relationship between IBA-1, GFAP and the alterations on the psychiatric behaviors by long term inhalation of combustible cigarette or e-cigarette, we analyzed the correlation between the expression levels of IBA-1 and GFAP and behavioral parameters. The results showed that there was a significant negative correlation between the expression of IBA-1 in LC brain and latency to nest in VLT (Pearson r = −0.5772, *p* = 0.0494, Fig. 7b). There was a significant negative correlation between the expression of GFAP in LC brain region and open arm time in EPM (Pearson r = −0.6349, *p* = 0.0265, Fig. 7c). According to the comprehensive analysis of the experimental results, the neuroimmunity of mice after long-term inhalation of cigarettes and aerosols is closely related to the brain region of LC.

**Figure 7.**
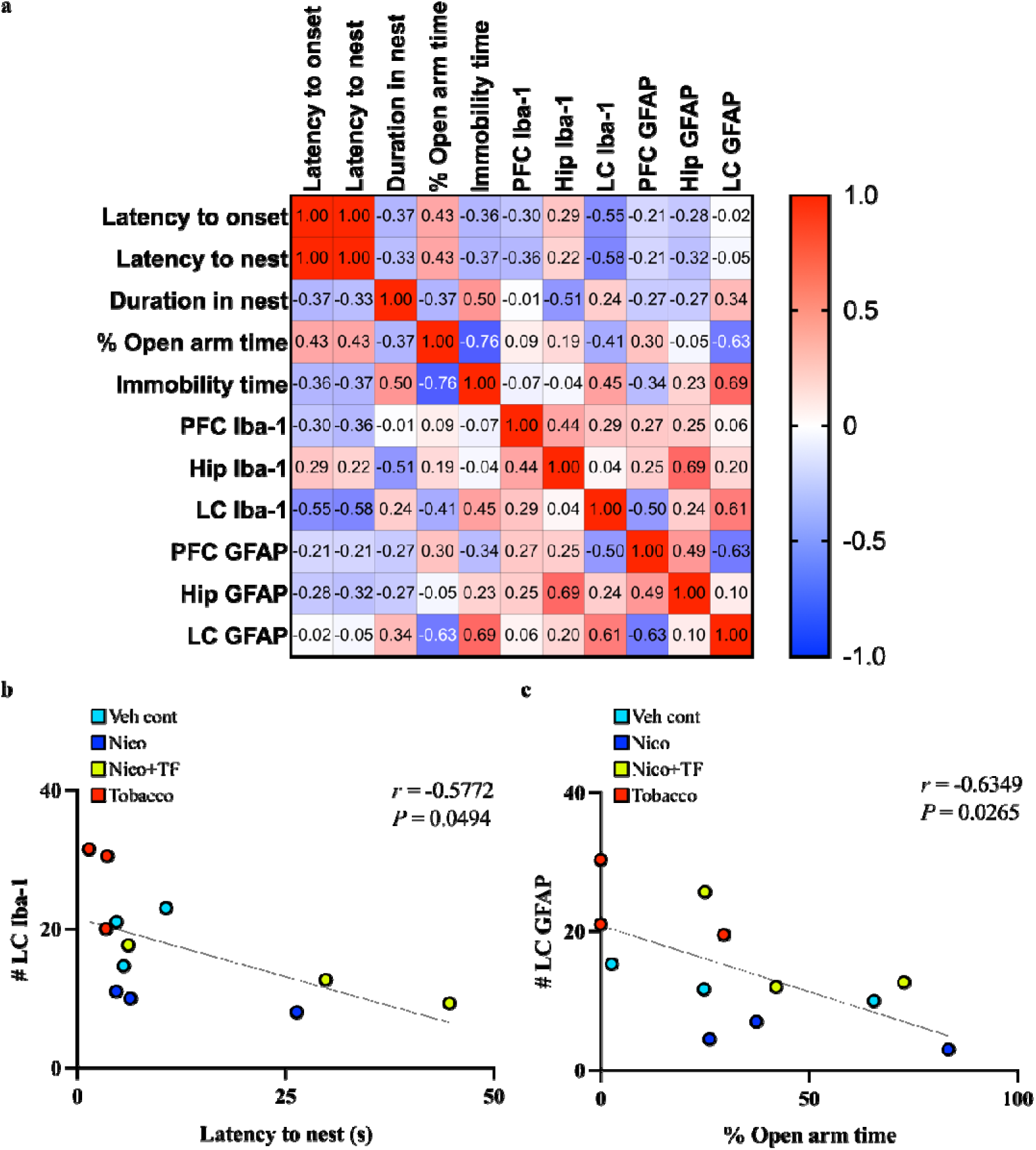
Correlation of glial cell expressions in LC region and behaviors. (a) Correlation diagram of glial cell expression in LC and behaviors. (b) Correlation between IBA-1 and latency to nest in VLT. (c) Correlation between GFAP and open arm time in EPM. Note: Veh cont, Vehicle control; Nico, Nicotine; Nico+TF, Nicotine with tobacco flavoring; Tobacco, Hongtashan Classic 1956 Hard Cigarette.

## Discussion

Smoking caused approximately 8 million deaths in 2019 and the global number of smokers continues to rise [1]. Despite their widespread use, there is currently insufficient data to determine if e-cigarettes are indeed safer than conventional tobacco cigarettes or useful as a smoking cessation aid. To prevent unforeseen outcomes like increasing nicotine use among young people and a future wave of diseases and fatalities linked to the use of nicotine delivery products, it is crucial to understand the neuropsychiatric effects of e-cigarettes as a new alternative nicotine tobacco product. Most e-cigarette products on the market provide a wide selection of flavoring agents combined with nicotine [40, 41], and there have been numerous toxicological and *in vivo* biological impact investigations on different flavors in electronic nicotine delivery devices conducted [28, 42–44], while the comprehensive understanding on the psychiatric effects of flavored e-vapors are currently very limited. The Food and Drug Administration (FDA) first approved e-cigarettes in 2021 but only permitted the flavors of tobacco [45]. Therefore in our present work, we exposed the mice to either the e-cigarette without flavors/with tobacco flavor, or combustible cigarette for one hour per day and lasted for more than a month. Our current data present the first evidence by CPP test revealing the aversive preference for the combustible cigarette in mice. The emotion, memory and social related-behavioral assessments indicated that the chronic exposure of combustible cigarette induced anxiety-like behaviors not only in the withdrawal period but also during vapor exposure, as assessed by the reduced latency to nest as the visually evoked innate fear response at the vapor exposure period; the elevated plus maze test during the vapor withdrawal stage also suggested that the significantly increased level of anxiety after exposure of the combustible cigarette, which were not found in mice exposed to e-cigarette with or without tobacco flavor. Nicotine was shown to activate a subpopulation of dopaminergic neurons in the ventral tegmental area (VTA) to induce anxiety [46], and in our present study, we have exposed the mice with aerosols or smoke containing the same concentration of nicotine and strikingly found the distinct psychiatric effects of combustible cigarette compared with e-cigarettes, even with the same flavor. Our data indicated that the route of administration of nicotine has critical roles on psychiatric behaviors, which may be more related to the formation temperature of the aerosol and the size of the particles, other than the flavors.

Despite the controversy, cigarette smoking and e-cigarette use have been confirmed to impact both innate and adaptive immune responses and regulate multisystem inflammatory response [35, 47]. The diverse ways of immune response triggered by nicotine which are manifested by either enhancing the immune reactions, or decreasing systemic activity against inflammation, represent the complexity of roles of nicotine on immunity. Here we presented and compared the cytokine levels including IL-6, IL-10, IL-17A, IL-1β, TNF-α and IFN-γ in the whole brain of mice chronically exposed to aerosol of combustible cigarette or e-cigarettes respectively, we have observed that IL-17A and IFN-γ were significantly decreased, while IL-1β and TNF-α were significantly increased by combustible cigarette, indicating that cigarette smoke exposure not only caused anxiety, but also increased the content of pro-inflammatory factors in the brain. Our data suggested that the actions on the psychiatric behaviors and neuroimmune profile are diverse among nicotine vapor with or without tobacco flavor, as well as combustible cigarettes in mice. Therefore, the mechanisms of action of tobacco cigarettes, e-cigarettes with or without additives could be complex.

Both microglia and astrocytes play vital roles in neuroinflammation and dysfunction of either cell type is associated with neuropsychiatric disorders [48]. Astrocyte dysfunction due to chronic stress or trauma can lead to the release of pro-inflammatory cells, loss of white matter, inhibition of axonal regeneration, formation of glial scarring, and inhibition of neurogenesis [49]. Microglia activation can lead to chronic inflammation, neuroinflammation and oxidative stress, and neurodegeneration [50]. Previous work has shown that nicotine acts in both directions, promoting and inhibiting inflammation [35]. Both chronic nicotine and withdrawal induce activation or remodeling of the highly adaptable resting microglia in the nucleus accumbens [51]. But it has also been reported that nicotine can act as an inhibit inflammation by activating α4β2-nAChRs [52]. Studies have shown that after neuronal injury, reactive astrocytes can promote neuronal destruction by synthesizing and releasing pro-inflammatory cytokines [53]. Reactive astrocytes can also actively promote secondary degeneration of the central nervous system after injury or in response to inflammatory signals [54]. It has been reported that primary culture of astrocytes from newborn rats exposed to cigarette smoke throughout gestation and incubated with hydrogen peroxide for 1 h shown significantly reduced antioxidant levels and cell survival [55]. As astrocytes are integral in maintaining central nervous system (CNS) homeostasis, astroglial cell loss could result in impairment of the antioxidant response leading to oxidative stress in the brain. The nitroamine (4-N-methyl-N-nitrosamino-1-(3-pyridyl)-1-butanone) in tobacco smoke causes inflammatory changes in the brain and neuronal damage by activating resident microglia and astrocytes [56].

To explore the levels of glial cells in specific brain regions that regulated by tobacco or e-cigarette aerosols, we performed immunofluorescent staining on the brain regions involved in neuropsychiatric behaviors especially the innate fear associated anxiety, and our data showed that the expression levels of IBA-1 and GFAP in LC brain area of mice in Tobacco group were significantly higher than that in Nico + TF group and Nico group, which indicated that long-term tobacco exposure could induce the elevated expression of microglia and astrocytes in LC brain region. This result is consistent with previous study, that is, nicoitine has potential neuroprotective effect, adding flavor to e-cigarette might not cause further neuroinflammation [24]. In VTA, the expression of IBA-1 in Tobacco group was significantly higher than that in Nico and Nico + TF groups, while the expression of GFAP in Nico group was significantly lower than that in Nico + TF and Tobacco groups. In HIP, the expression of IBA-1 had no significant difference, but the expression of GFAP in Nico + TF group was significantly lower than that in Nico and Tobacco groups. In SNr, the expression of IBA-1 and GFAP in Tobacco group was significantly higher than that in Nico group. There was no significant difference in the expression of IBA-1 in PFC, but the expression of GFAP in Nico group increased significantly. There was no significant difference in the expression of IBA-1 and GFAP in BLA. However, the mechanism of nicotine in these emotion-related brain regions has not been reported. Through the linear correlation analyses between the behavioral parameters of mice and the levels of IBA-1 and GFAP, we have observed the significantly negative correlations between the IBA-1 level in LC and the latency to nest in VLT; as well as the GFAP level in LC and the open arm time in EPM. LC is one of the key brain nuclei implicated in innate fear as well as the pathophysiology of anxiety and depression [57–59] and was shown activated by nicotine [60], our data may indicate that the chronic cigarette inhalation induces anxiety via the dramatic effects on the neuroimmune functions in the LC region.

## Conclusion

Our findings suggest that chronic aerosol inhalation with combustible cigarette induced significantly higher levels of innate fear and anxiety behaviors compared to e-cigarettes, which was shown unrelated to the tobacco flavor. Combined with previous evidence that e-cigarette expose increased neurologic risk, the long-term aerosol exposures to the nicotine delivery device including combustible cigarettes and e-cigarettes, lead to alterations on the neuroimmune profile in mice, especially the levels of microglia and astrocytes in the locus coeruleus were shown a negatively linear correlation with innate fear and anxiety, suggesting the glial signals in LC could be a therapeutic target for smoking-related anxiety. Further research is greatly needed to better understand the different neurological impacts of tobacco cigarettes and e-cigarettes including the neural circuit underlying nicotine induced chronic stress via various delivery methods, to develop better strategies for smoking related anxiety.

## Supplemental data

**Figure S1.**
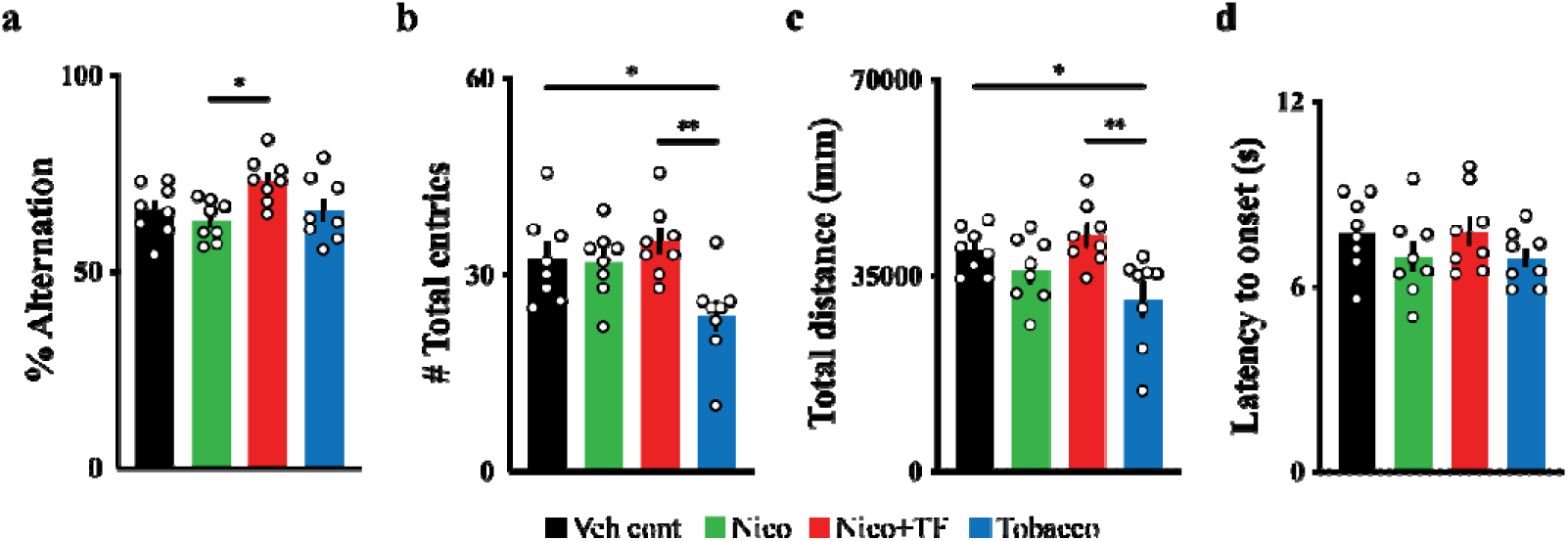
Spatial memory assessment (a-c). The percentage alternation was calculated by dividing spontaneous alternation by (the total number of alternations minus two). (d) Perceptual behavioral assessment. The latency to overt behavioral sign of nociception. \**p* < 0.05 and \*\**p* < 0.01 as determined by Ordinary one-way ANOVA, multiple comparisons with every other group. Bars represent marginal means ± SEM. N=8 per group. Note: Veh cont, Vehicle control; Nico, Nicotine; Nico+TF, Nicotine with tobacco flavoring; Tobacco, Hongtashan Classic 1956 Hard Cigarette. EPM, Elevated plus maze.

**Table S1.** Nicotine concentration of all groups.

| Drug | Nicotine content (g/kg) |
| --- | --- |
| Vehicle control | 0 |
| Nicotine | 40 |
| Nicotine with tobacco flavoring | 40 |
| Tobacco cigarette | 40 |

## Author Contributions

Conceptualization, Zuxin Chen and Xin-an Liu; Data curation, Jiayan Ren, Zixuan Li, Ruoxi Wang, Zehong Wu, Xin-tao Jiang, Liping Wang, Zuxin Chen and Xin-an Liu; Funding acquisition, Xin-an Liu; Investigation, Zhibin Xu, Jiayan Ren, Xiaoyuan Jing, Zhi-zhun Mo, Zixuan Li, Yiqing Zhao, Ruoxi Wang and Ye Tian; Methodology, Zhibin Xu, Jiayan Ren, Xiaoyuan Jing, Zhi-zhun Mo and Zehong Wu; Project administration, Xin-an Liu; Resources, Liping Wang and Xin-an Liu; Supervision, Zuxin Chen; Validation, Zhibin Xu, Xiaoyuan Jing and Xin-tao Jiang; Writing – original draft, Zhibin Xu, Xiaoyuan Jing and Xin-an Liu; Writing – review & editing, Zuxin Chen and Xin-an Liu.

